# A comprehensive dataset of TLX1 positive ALL-SIL lymphoblasts and primary T-cell acute lymphoblastic leukemias

**DOI:** 10.1101/695668

**Authors:** Karen Verboom, Wouter Van Loocke, Emmanuelle Clappier, Jean Soulier, Jo Vandesompele, Frank Speleman, Kaat Durinck

## Abstract

Most currently available transcriptome data of T-cell acute lymphoblastic leukemia (T-ALL) are based on polyA[+] RNA sequencing methods thus lacking non-polyadenylated transcripts. Here, we present the data of polyA[+] and total RNA sequencing in the context of *in vitro TLX1* knockdown in ALL-SIL cells and a primary T-ALL cohort. We extended this dataset with ATAC sequencing and H3K4me1 and H3K4me3 ChIP sequencing data to map putative gene regulatory regions. In this data descriptor, we present a detailed report of how the data were generated and which bioinformatics analyses were performed. Through several technical validations, we showed that our sequencing data are of high quality and that our *in vitro TLX1* knockdown was successful. We also validated the quality of the ATAC and ChIP sequencing data and showed that ATAC and H3K4me3 ChIP peaks are enriched at transcription start sites. We believe that this comprehensive set of sequencing data can be reused by others to further unravel the complex biology of T-ALL in general and TLX1 in particular.

## Background & Summary

T-cell acute lymphoblastic leukemia (T-ALL) is a hematological cancer resulting from malignant transformation of normal precursor T-cells. T-ALL accounts for 15 % of the pediatric and 25 % of the adult ALL cases and can be subdivided into four molecular subgroups (TLX, HOXA, immature and TAL-R), each with a unique gene expression profile signature^1,2^. In addition, several other oncogenes (eg *NOTCH1*) and tumor suppressor genes (eg *CDKN2A* and *PTEN*), are involved in the multi-step process of leukemia formation across these subgroups^3,4^. Besides genetic alterations, also epigenetic mechanisms, such as deregulation of enhancers and histone modifications, are disturbed in T-ALL^5^.

Several sequencing efforts, including whole genome, exome and transcriptome sequencing, have been performed to identify driver genes in T-ALL. The majority of these studies focused on the identification of mutations, fusion genes and transcriptional changes^6–8^. A major drawback of the current transcriptome studies is that polyA[+] RNA-seq only covers part of the transcriptome and provides no insights into the non-polyadenylated complement that includes biologically relevant transcripts such as circular RNAs (circRNAs) and long non-coding RNAs (lncRNAs)^9,10^.

To resolve this issue, we generated polyA[+] as well as total RNA-seq data of 60 (17 TLX, 7 HOXA, 13 immature, 23 TAL-R) and 25 (10 TLX, 5 HOXA, 5 immature, 5 TAL-R) T-ALL patients, respectively. In this dataset, we identified TLX subgroup specific and possibly oncogenic lncRNAs as reviewed by Verboom et al.^11^. As we focused on the TLX subgroup and more specifically on TLX1, we also generated polyA[+] and total RNA-seq data of *TLX1* knockdown samples generated using two independent siRNAs in triplicate. In addition, we performed assay for transposase accessible chromatin sequencing (ATAC-seq) as well as H3K4me3 and H3Kme1 chromatin immunoprecipitation sequencing (ChIP-seq) and used the publicly available TLX1 ChIP data (GSE62264) to identify regions directly regulated by TLX1 in TLX1 positive ALL-SIL lymphoblasts. This is a unique dataset in the T-ALL field as it combines for the first time polyA[+] and total RNA-seq of an *in vitro* model system as well as a large primary T-ALL cohort. Moreover, this dataset was extended with ChIP-seq and ATAC-seq to define gene regulatory regions.

Here, we provide a detailed description of the methods and bioinformatics analyses used to generate the data to facilitate data-repurposing by other researchers. Moreover, we demonstrate that our data is of high quality. The T-ALL cohort has also been annotated according to their genetic subgroup. In our related publication, we used the dataset to identify TLX1 regulated lncRNAs and TLX subgroup specific lncRNAs^11^. Therefore, this dataset can be reused to characterize other subgroup specific or TLX1 regulated biotypes or to study other subgroups in the primary T-ALL cohort as this has only been done using microarray data^12^. Besides gene expression analysis, this RNA-seq dataset can also be used to identify gene mutations and gene fusions^6^. Furthermore, integrating the ChIP-seq and ATAC-seq data, together with GRO-seq data can be useful to study enhancer RNAs in TLX1 positive T-ALL.

In summary, this data descriptor provides detailed information about the methods used to generate and analyze ChIP-seq and ATAC-seq data in the TLX1 positive ALL-SIL cell line as well as transcriptome data through polyA[+] and total RNA-seq of a *TLX1* knockdown *in vitro* model system and a primary T-ALL cohort. As this is a rich dataset including transcriptome as well as epigenome data, we believe that this dataset is of great value to the research community and will aid, amongst others, to further unravel the complex biology of T-ALL.

## Methods

### Cell lines

The TLX1 positive cell line ALL-SIL was obtained from the DSMZ cell line repository (ACC 511). Cells were maintained in RPMI-1640 medium (Life Technologies, 52400-025) supplemented with 20 % fetal bovine serum (PAN Biotech, P30-3306), 1 % of L-glutamine (Life Technologies, 15140-148) and 1 % penicillin/streptomycin (Life Technologies, 15160-047) at 37 °C in a 5 % CO_2_ atmosphere. Short tandem repeat genotyping was used to validate cell line authenticity prior to performing the described experiments and mycoplasma testing is done on a monthly basis in our laboratory.

### siRNA mediated knockdown in ALL-SIL lymphoblasts

The RNA isolated upon *TLX1* knockdown that has been used for microarray based gene expression profiling in a previous study has been used in this study for RNA-seq^13^. In short, 400 nM of Silencer Select Negative Control 1 siRNA (Ambion, #AM4635) or siRNAs targeting TLX1 (Ambion, Carlsbad, #4392420, s6746 (siRNA1) and s6747 (siRNA2)) was added to 8 million ALL-SIL cells in a total volume of 400 μl. The samples were electroporated (250 V, 1000 μF) in electroporation cuvettes (Bio-Rad, 1652086) using a Genepulser Xcell device (Bio-Rad). ALL-SIL cells were collected 24h post electroporation.

### Clinical samples

Bone marrow lymphoblast and blood samples from 60 pediatric and adult T-ALL patients (13 immature, 23 TAL-R, 17 TLX1/TLX3 and 7 HOXA) were collected with informed consent according to the declaration of Helsinki from Saint-Louis Hospital (Paris, France) and the study was approved by the Institut Universitaire d’Hématologie Institutional Review Board and the Ghent University Hospital (approval number B670201627319). Subgroup annotation is given in Supplementary Table 1. White blood cells from the patients samples were isolated by Ficoll centrifugation and cryopreserved using standard procedures. After thawing, cells were spun down and RNA was extracted^14^.

### RNA isolation

Total RNA was isolated using the miRNeasy mini kit (Qiagen, 217084) with DNA digestion on-column according to the manufacturer’s instructions. By means of spectrophotometry, RNA concentrations were measured (Nanodrop 1000, Thermo Scientific). RNA quality scores (RNA integrity number (RIN)) of the RNA of the cell line experiments were high (RIN: 9-10). The RIN score of the RNA of the primary samples were determined using a bioanalyzer and are shown in Supplementary Table 1. All bioanalyzer profiles suggest RNA of good quality.

### PolyA[+] RNA sequencing

Library prep of the polyA[+] transcripts of the *TLX1* knockdown samples was performed using the TruSeq stranded mRNA sample preparation kit (Illumina, RS-122-2101) according to manufacturer’s instructions. The libraries were quantified using the KAPA library quantification kit (Roche, KK4854) and samples were paired-end sequenced on the NextSeq 500 sequencer (Illumina) with a read length of 2 × 75 bp. PolyA[+] RNA-seq of the 60 primary T-ALL samples was performed using 100 ng of RNA as input material by Biogazelle (Belgium) with the TruSeq stranded mRNA sample preparation kit according to manufacturer’s instructions. The samples were quantified using the KAPA library quantification kit and paired-end sequenced on a NextSeq 500 sequencer with a read length of 2 × 75 bp. The number of uniquely mapped reads are shown in Supplementary Table 1-2.

### Total RNA sequencing

Total RNA-seq of the *TLX1* knockdown samples and 25 primary T-ALL samples (5 immature, 5 TAL-R, 10 *TLX1*/*TLX3* and 5 HOXA) was performed using the TruSeq Stranded Total RNA (w/RiboZero Gold) sample prep kit (Illumina, RS-122-2301), involving depletion of ribosomal (rRNA) transcripts by Biogazelle according to manufacturer’s instructions. Libraries were quantified using the Qubit 2.0 Fluorometer (Thermo Fischer Scientific) and paired-end sequenced on a NextSeq 500 sequencer with a read length of 2 × 75 bp. The number of uniquely mapped reads are shown in Supplementary Table 1-2.

### Data processing RNA sequencing

FastQC v0.11.3 was used for quality control of fastq files (available online at http://www.bioinformatics.babraham.ac.uk/projects/fastqc). All samples were aligned against the human reference genome (GRCh38) with STAR_2.4.2a^15^ using default settings and two pass methods as described in the STAR manual. GENCODE v25 primary annotation was used as a guide during the first pass and the combined splice junction database of all samples generated in the first pass as a guide in the second step. Genes were quantified on GENCODE v25 during alignment with STAR. Raw data files have been deposited in the NCBI Gene Expression Omnibus (GEO) database^16-19^. MultiQC v0.9 was used to aggregate the FastQC and STAR results of all samples. Sample IDs and subgroups are shown in Supplementary Table 1. TDF files were loaded in IGV v2.3.98 to visualize the data.

### Assay for transposase accessible chromatin sequencing

Assay for transposase accessible chromatin sequencing (ATAC-seq) was performed as previously described with minor changes^20^. In short, 50,000 cells were lysed and fragmented using Tn5 transposase (Illumina, FC-121-1030). Next, the samples were purified using the MinElute kit (Qiagen, 28204). Ten μl transposased DNA fragments were amplified by adding 10 μl H_2_O, 2.5 μl of each forward (5’AATGATACGGCGACCACCGAGATCTACACTCGTCGGCAGCGTCAGATGTG3’) and reverse (5’CAAGCAGAAGACGGCATACGAGATAAAATGGTCTCGTGGGCTCGGAGATGT3’) primer (25 μM) and 25 μl NEBNext High-Fidelity 2x PCR master mix (Bioké, M0541) using the following PCR program: 5 min at 72 °C, 30 sec at 98 °C and 5 cycles of 10 sec at 98 °C, 30 sec at 63 °C and 1 min at 72 °C. To reduce GC-content and size bias, qPCR was performed to determine the exact number of PCR cycles to prevent saturation. 5 μl DNA, 3 μl H_2_O, 1 μl of each forward and reverse primer (5 uM) and 10 μl SYBR green (Roche, 4707516001) were used for qPCR (30 sec at 98 °C and 19 cycles of 10 sec at 98 °C, 30 sec at 63 °C and 1 min at 72 °C). The number of extra PCR cycles was calculated as the number of qPCR cycles that correspond to ¼ of the maximum fluorescent intensity. Ten extra PCR cycles were performed on the 45 μl transposed DNA fragments. Samples were purified using the PCR Cleanup kit (Qiagen, 28104). Sample quality was determined using Fragment Analyzer (Advanced Analytical) and samples were quantified using the KAPA library quantification kit. Samples were single-end sequenced on a NextSeq 500 with a read length of 75 bp and an average sequencing read depth of 81.5 million reads.

### Chromatin immunoprecipitation sequencing

H3K4me1 and H3K4me3 chromatin immunoprecipitation sequencing (ChIP-seq) were performed as previously described with minor changes^21^. In brief, 1×10^7^ cells were cross-linked with 1.1 % formaldehyde (Sigma-Aldrich, F1635) at room temperature for 10 min and the cross-linking reaction was quenched with an access of glycine (125 mM final concentration, Sigma-Aldrich, G-8790). Nuclei were isolated and chromatin was purified by chemical lysis. Next, the purified chromatin was fragmented to 200-300 bp fragments by sonication (Covaris, M220, Focused-ultrasonicator). Chromatin immunoprecipitation was performed by incubation of the chromatin fraction overnight with 20 μl of protein-A coated beads (Thermo-Scientific, 53139) and 2 μg of H3K4me1 specific (Abcam, ab8895) or 2 μg of H3K4me3 specific (Abcam, ab8580) antibody. The next day, beads were washed to remove non-specific binding events and enriched chromatin fragments were eluted from the beads, followed by reverse cross-linking by incubation at 65 °C overnight. DNA was subsequently purified by phenol/chloroform extraction, assisted by phase lock gel tubes (5Prime, 733-2478). DNA obtained from the ChIP-assays was adaptor-ligated and amplified using the NEBNext multiplex oligo kit (New England BioLabs, E7335S) according to manufacturer’s instructions. Libraries were quantified using the KAPA library quantification kit. The input, H3K4me1 and H3K4me3 libraries were single-end sequenced on a NextSeq 500 sequencer with a read length of 75 bp and an average sequencing depth of 32.9 million reads.

### Data processing ChIP-seq and ATAC-seq

FastQC was used for quality control of fastq files. All ChIP-seq and ATAC-seq files were aligned against the human reference genome (GRCh38) using STAR_2.4.2a^15^ with -- outFilterMultimapNmax 1 to exclude multimapping reads and --alignIntronMax 1 to turn off splice awareness. MultiQC was used to aggregate the FastQC and STAR results of all samples. Raw data files have been deposited in the NCBI Gene Expression Omnibus (GEO) database^22-23^. To detect enriched regions (ChIP-seq) and regions of open chromatin (ATAC-seq), peak calling has been performed with MACS2 v2.1.20150731 using input samples as control for ChIP-seq^24^. Peak calling for histone marks was performed using --broad setting. Generated files show peak location, enrichment of the peak and score of the identified peaks. TDF and bed files were loaded in IGV v2.3.98 to visualize the data. ChIPseeker v1.16.1 was used to identify enrichment of ATAC-seq peaks around the transcription start sites^25^.

### Code availability

Quality control and data analyses were performed with following software and versions, as described in the method section:

1. FastQC, version v0.11.3 was used for the quality analysis of the raw fastq files of the RNA-seq, ChIP-seq and ATAC-seq data. (http://www.bioinformatics.babraham.ac.uk/projects/fastqc)
2. For RNA-seq, STAR v2.4.2a was used for mapping the reads to the Hg38 human reference genome using default settings and two pass methods as described in the STAR manual. GENCODE v25 primary annotation was used as a guide during the first pass and the combined splice junction database of all samples generated in the first pass as a guide in the second step. Genes were quantified on GENCODE v25 during alignment with STAR. For ATAC-seq and ChIP-seq, STAR v2.4.2 was used for mapping the reads to the Hg38 human reference genome with --outFilterMultimapNmax 1 to exclude multimapping reads and --alignIntronMax 1 to turn off splice awareness.
3. FastQC and STAR results were aggregated using MultiQC v0.9.
4. Peak calling for the ATAC-seq and ChIP-seq data was performed with MACS2 v2.1.20150731 using input samples as control for ChIP-seq. Peak calling for the histone marks was performed using --broad setting.

## Data Records

Sample IDs and subgroup annotation are shown in Supplementary Table 1-2. For RNA-seq, the raw data files and count tables have been deposited in the GEO database. For polyA[+] RNA-seq of the *TLX1* knockdown samples, the accession number is GSE110632 (2018)^16^; for total RNA-seq of the *TLX1* knockdown samples, the accession number is GSE110635 (2018)^17^; for polyA[+] RNA-seq of the primary T-ALL cohort, the accession number is GSE110633 (2018)^18^; for total RNA-seq of the primary T-ALL cohort, the accession number is GSE110636 (2018)^19^; for ATAC-seq of the ALL-SIL cell line, the accession number is GSE110630 (2018)^22^; for ChIP-seq data, the accession number is GSE110631 (2018)^23^. These datasets were submitted as SubSeries of the SuperSeries GSE110637 (2018).

## Technical Validation

### Validation of RNA sequencing data

To validate the quality of our RNA-seq data, FastQC analyses were performed on the fastq files. The mean quality scores for the polyA[+] and total RNA-seq data of the *TLX1* knockdown samples and the primary T-ALL cohort are shown in Figure 1a. These plots show the mean quality score across each base position per sample and indicate by a color scale if the quality is good (green), reasonable (orange) or bad (red). As the mean quality score per base position is located in the “good quality region” for each sample, we confirm that our sequencing data is of high quality. Of note, a small decrease in the quality towards the end of a read is typically observed, which is inherent to the Illumina sequencing by synthesis procedure^26^. As these samples have a high sequencing coverage, further increasing the number of reads would only give a small increase in the number of detectable genes (Fig. 1b). A high percentage of the reads were uniquely mapped to the Hg38 human reference genome, confirming a good mapping quality (Supplementary Table 1-2, Fig. 1b). After mapping, we detected an average of 26,412 genes over all samples.

**Figure 1:**
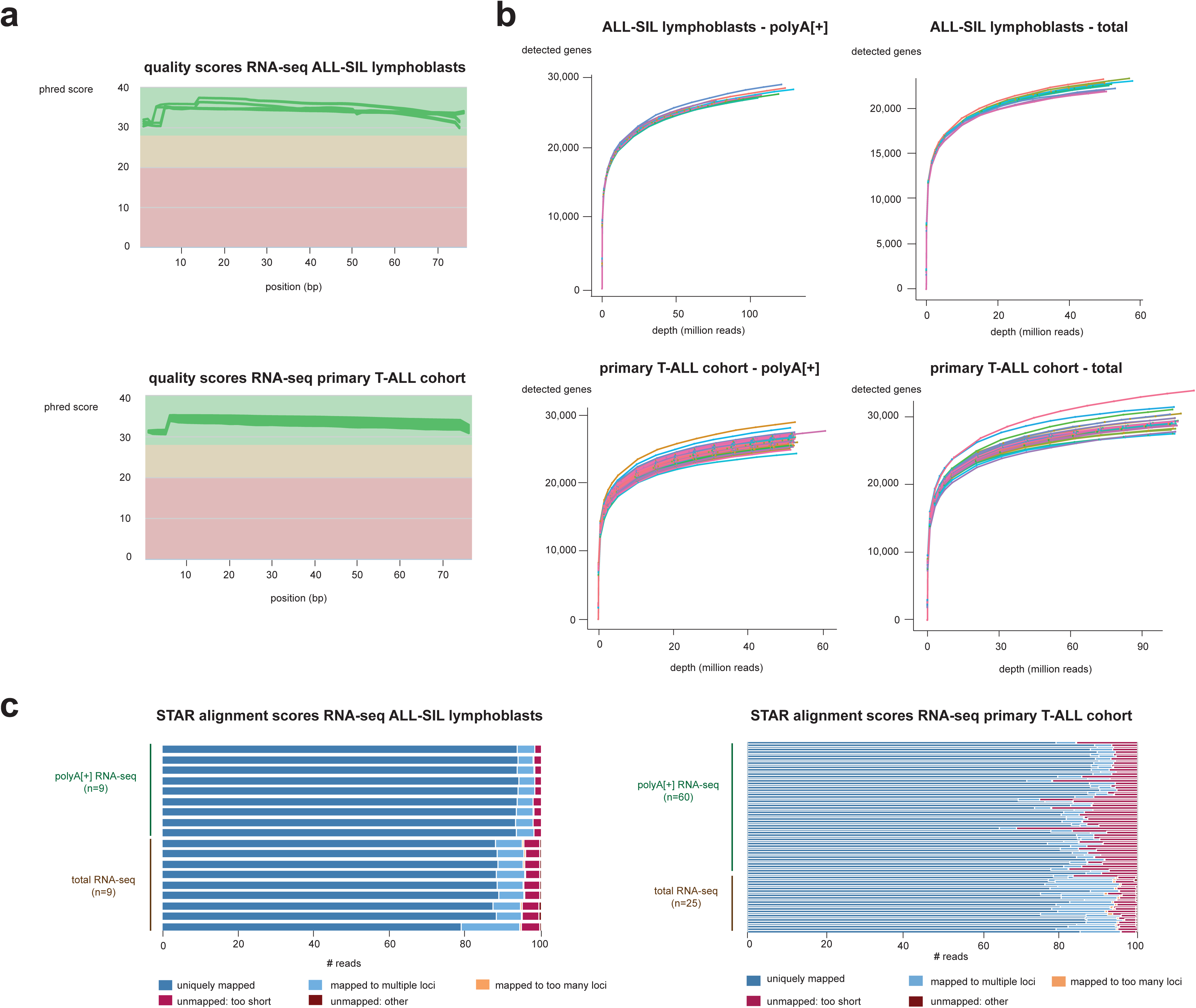
Quality assessment of polyA[+] and total RNA-seq data of the *TLX1* knockdown and primary T-ALL cohort samples. (a) Average quality score per base position per sample generated by combining FastQC and MultiQC for the *TLX1* knockdown samples (upper panel) and the primary T-ALL cohort (lower panel). Each line represents a sample. Scores greater than 30 (green region) indicate a good quality. (b) Sequencing saturation plots showing the number of genes detected at a given sequencing depth. (c) Barplots showing the STAR alignment scores for the *TLX1* knockdown samples (left panel) and the primary T-ALL cohort (right panel) generated with MultiQC. Black dots show the average expression.

### Validation of *TLX1* knockdown in ALL-SIL lymphoblasts

Knockdown of *TLX1* has been validated as *TLX1* is significantly lower expressed in the knockdown samples as compared to the control samples in the three replicates of polyA[+] (66 %, 58 % and 65 % knockdown) and total RNA-seq (56 %, 48 % and 62 % knockdown) data (Fig. 2a). In addition, we could demonstrate that the well-known tumor suppressor genes *FAT1* and *PTPN2*, shown to be repressed by TLX1 by microarray data^13^, are significantly upregulated upon *TLX1* knockdown according to polyA[+] and total RNA-seq data (Fig. 2 b-c). Furthermore, correlation analysis shows that the replicates are concordant, as shown for a representative example (Fig. 2d).

**Figure 2:**
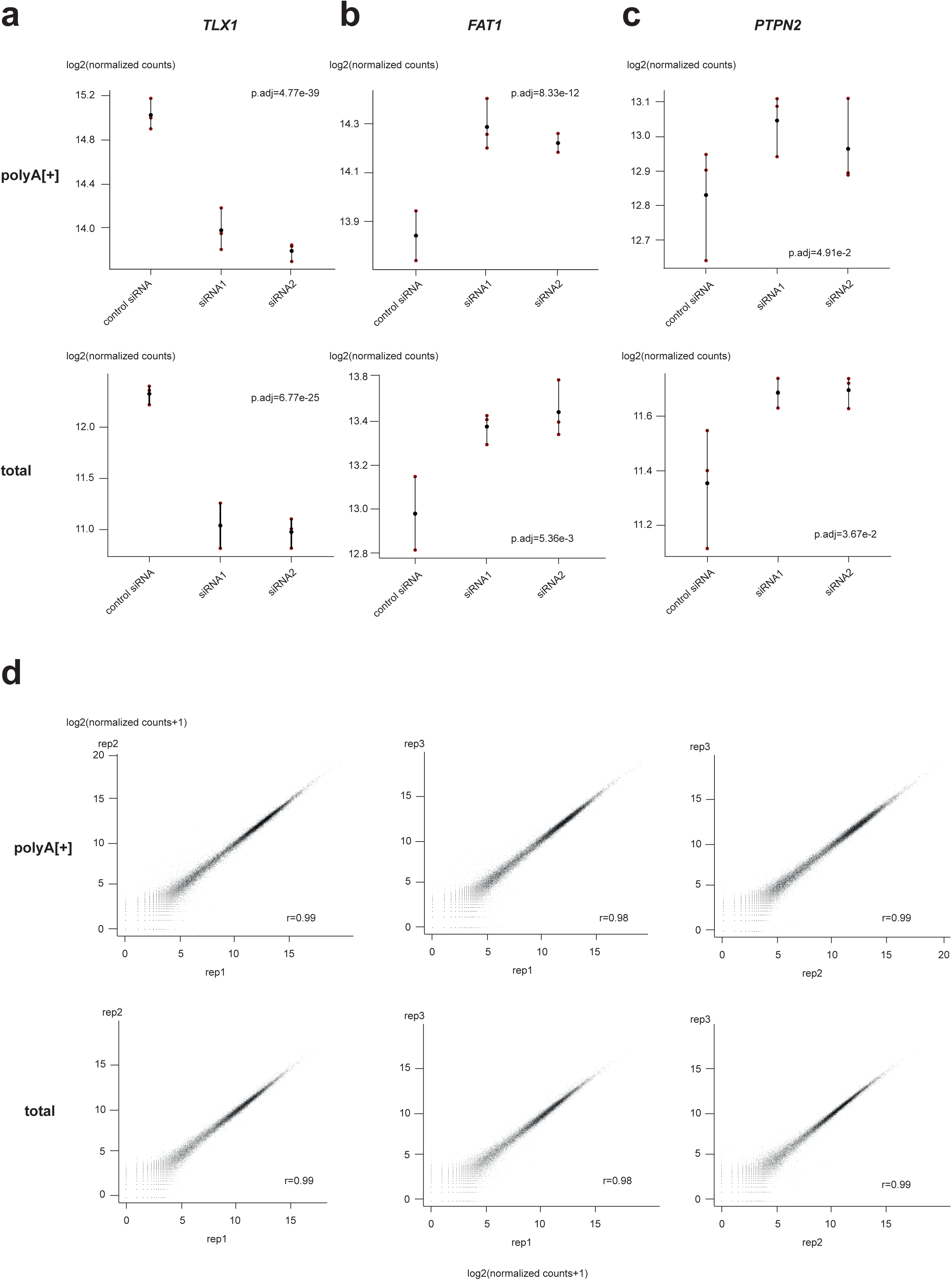
Validation of *TLX1* knockdown in ALL-SIL lymphoblasts. (a) *TLX1* is significantly downregulated in the *TLX1* knockdown samples. Tumor suppressor genes *FAT1* (b) and *PTPN2* (c), known to be repressed by *TLX1*, are upregulated upon *TLX1* knockdown. (d) correlation plots for the replicates of siRNA 1. (d) Correlation plots for the three replicates of TLX1 siRNA1 for polyA[+] (upper panel) and total RNA-seq (lower panel). Pearson correlation coefficients are shown.

### Validation of ATAC sequencing data

Before sequencing, we validated the quality of the ATAC-seq library by inspecting Fragment Analyzer profiles. As expected for ATAC-seq libraries, the profile first showed a high peak around 180 nt containing the nucleosomal free DNA fragments, followed by two peaks of fragments containing 1 or 2 nucleosomes, respectively (Fig. 3a). After sequencing, the quality of the raw sequencing data was validated by performing FastQC. The average base quality score fell in the “good quality region”, confirming that we generated high quality ATAC-seq data (Fig. 3b). Raw sequencing reads were aligned against the Hg38 human reference genome resulting in 81.5 million mapped reads. 76.27 % of the reads mapped uniquely to the human reference genome and were used for subsequent analyses (Fig. 3c). In addition, we also confirmed that there is an enrichment of ATAC peaks around transcription start sites, known to contain open chromatin (Fig. 3d).

**Figure 3:**
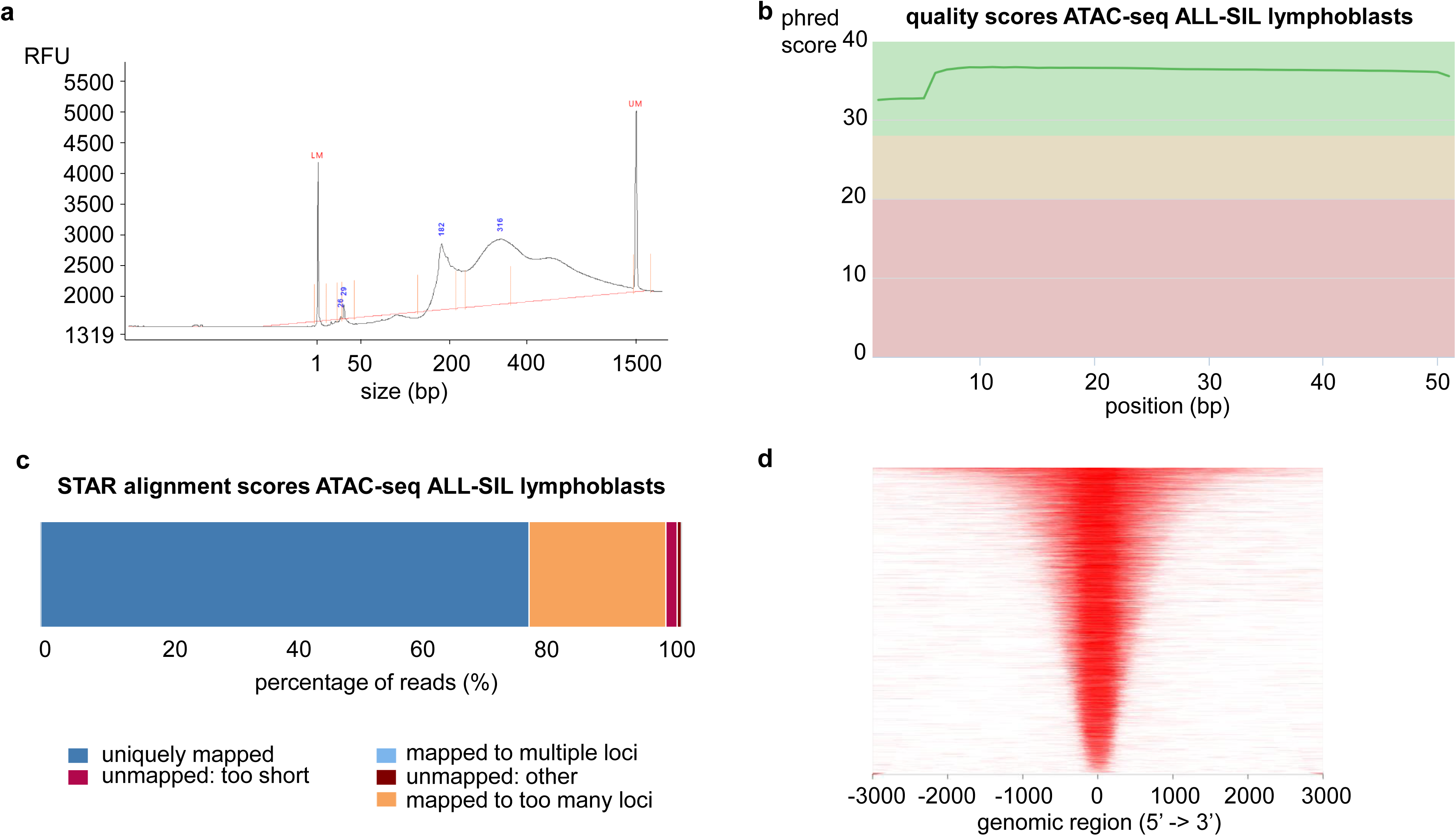
Quality assessment of ATAC-seq in ALL-SIL lymphoblasts. (a) Fragment analyzer profile showing a high nucleosome-free peak followed by a mono-nucleosome and dinucleosome peak. (b) Average quality score per base position generated by combining FastQC and MultiQC. Scores greater than 30 (green region) indicate a good quality. (c) Barplot showing the STAR alignment scores. (d) Enrichment of ATAC peaks around transcription start sites.

### Validation of ChIP-seq data

The quality of the raw sequencing data was determined using FastQC analyses and we could show that the average base quality score fell in the “good quality region” for each ChIP-seq sample (Fig. 4a). On average, 32.9 million reads per sample were aligned to the Hg38 human reference genome. 69.79 %, 89.49 % and 89.85 % of the reads were uniquely aligned to the Hg38 human reference genome for the input, H3K4me1 and H3K4me3 sample, respectively (Fig. 4b). As H3K4me3 is a marker of promotor regions, we showed that H3K4me3 is enriched on the promotor of *EEF2*, one of the highest expressed genes in parental ALL-SIL lymphoblasts (Fig. 4c).

**Figure 4:**
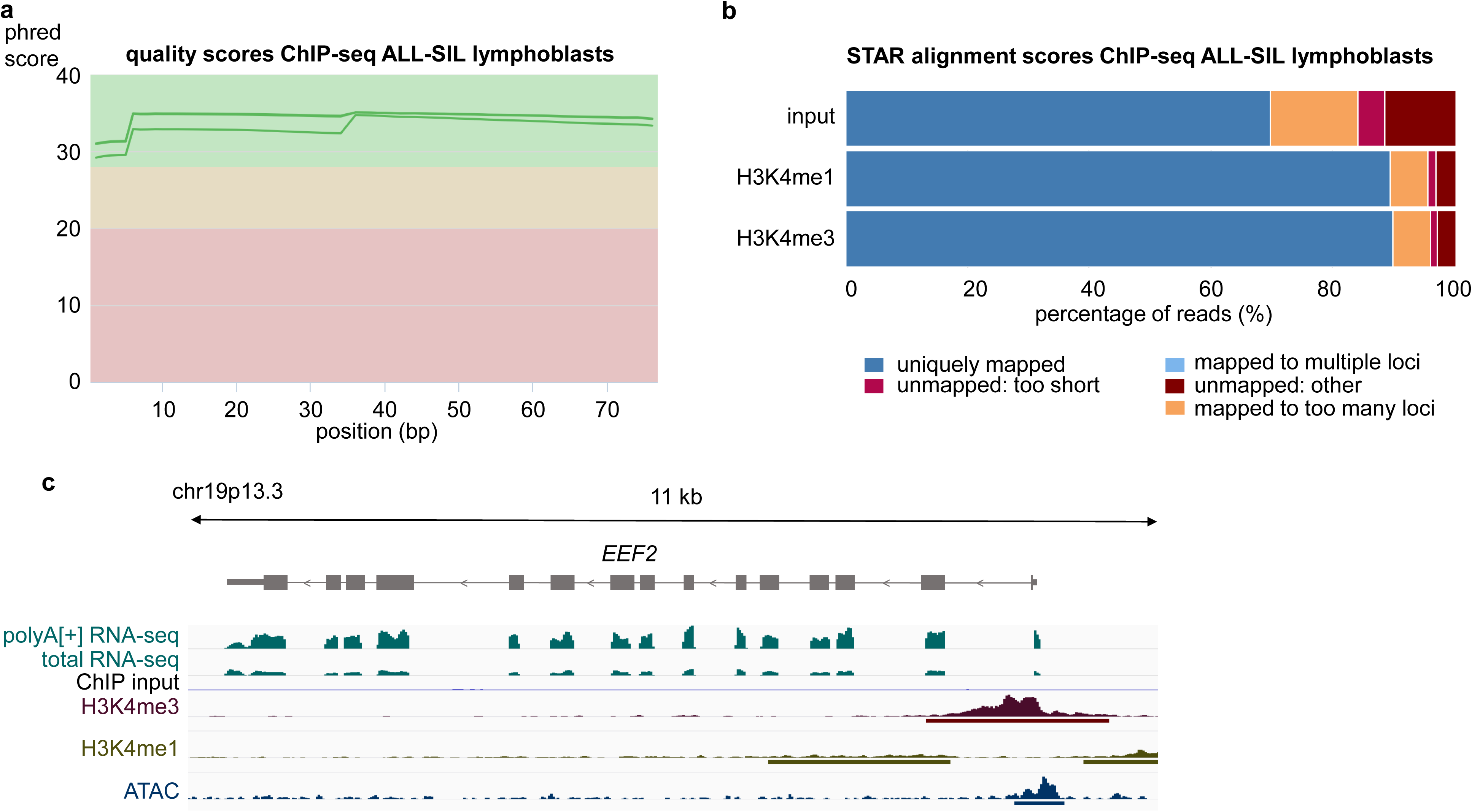
Quality assessment of H3K4me1 and H3K4me3 ChIP-seq in ALL-SIL lymphoblasts. (a) Average quality score per base position generated by combining FastQC and multiQC. Scores greater than 30 (green region) indicate a good quality. (BbBarplots showing the STAR alignment scores per sample. (c) IGV screenshot showing H3K4me3, H3K4me1, ChIP input and ATAC-seq peaks on the promotor of *EEF2*, one of the highest expressed genes in parental ALL-SIL lymphoblasts. PolyA[+] and total RNA-seq tracks are depicted for control siRNA transfected samples. ATAC, H3K4me3 and H3K4me1 bars represent the MACS2 peak with FDR < 0.05.

In conclusion, this data descriptor shows that we have generated high quality RNA-seq, ChIP-seq and ATAC-seq data that can be reused by other users to further unravel the complex biology of T-ALL.

### Usage Notes

The raw data and gene count tables of the polyA[+] and total RNA-seq data can be downloaded from the GEO database via accession numbers GSE110632^16^ (polyA[+]) and GSE11065^17^ (total) for the *TLX1* knockdown samples and via GSE110633^18^ (polyA[+]) and GSE110636^19^ (total) for the primary T-ALL cohort. The sample IDs and subgroup annotation can be found in Supplementary Table 1. The raw data and peak tables of the ATAC-seq and ChIP-seq can also be downloaded from the GEO database via accession numbers GSE110631^22^ (ATAC-seq) and GSE110630^23^ (ChIP-seq). The analyses can easily be repeated using the specifications described in the method section starting from the raw sequencing data. By providing the raw sequencing data, users can also use their own pipeline to analyse the data. Other pipelines, such as the bcbio-nextgen pipeline (https://github.com/bcbio/bcbio-nextgen), can also be used to analyze the ChIP-seq and ATAC-seq data. Matching TLX1 and H3K27ac ChIP-seq data in the ALL-SIL cell line, generated by Durinck et al., can be integrated with the datasets described in this paper and are available through GEO (GSE70734, GSE62264)^13^. Downstream differential analysis can be easily performed using the provided count tables and sample annotation (Supplementary Table 1-2) as input for Deseq2^27^ (as described by Verboom et al. ^11^) or other packages as Limma^28^ or EdgeR^29^.

## Acknowledgements

The authors want to thank the following funding agencies: Kinderkankerfonds, the Fund for Scientific Research Flanders (FWO Vlaanderen, research project G051416N; postdoctoral grant to KD), BOF (PhD grant to KV), Ghent University (GOA grant BOF16/GOA/023) and ANR 10-IBHU-0002 to JS. The computational resources (Stevin Supercomputer Infrastructure) and services used in this work were provided by the VSC (Flemish Supercomputer Center), funded by Ghent University, FWO and the Flemish Government – department EWI.

## Author contributions

Karen Verboom performed laboratory experiments and data analysis, was involved in the design of the experiments, coordination of the research, generation of the figures and writing of the manuscript.

Wouter Van Loocke assisted with the data analysis of RNA-seq, ChIP-seq and ATAC-seq. Jean Soulier and Emmanuelle Clappier collected the primary T-ALL samples.

Jo Vandesompele coordinated the research, designed experiments and helped writing the paper.

Frank Speleman coordinated the research, designed experiments and helped writing the paper.

Kaat Durinck coordinated the laboratory experiments, was involved in the design of the experiments and coordination of the research and helped writing the paper.

All authors approved the final version of the manuscript.

## Competing interests

There are no conflicts of interest.

## Table Legends

**Supplementary Table 1**: sample information of the primary T-ALL cohort.

**Supplementary Table 2**: sample information of ALL-SIL lymphoblasts.

